# Brain-wide electrical spatiotemporal dynamics encode reward anticipation

**DOI:** 10.1101/813113

**Authors:** Mai-Anh T. Vu, Lisa K. David, Gwenaëlle E. Thomas, Meghana Vagwala, Caley Burrus, Neil M. Gallagher, Joyce Wang, Cameron Blount, Dalton N. Hughes, Elise Adamson, Nkemdilim Ndubuizu, Il Hwan Kim, Scott Soderling, Stephen D. Mague, R. Alison Adcock, Kafui Dzirasa

## Abstract

Anticipation of an upcoming stimulus induces neural activity across cortical and subcortical regions and influences subsequent behavior. Nevertheless, the network mechanism whereby the brain integrates this information to signal the anticipation of rewards remains relatively unexplored. Here we employ multi-circuit electrical recordings from six brain regions as mice perform a sample-to-match task in which reward anticipation is operationalized as their progress towards obtaining a potential reward. We then use machine learning to discover the naturally occurring network patterns that integrate this neural activity across timescales. Only one of the networks that we uncovered signals responses linked to reward anticipation, specifically relative proximity and reward magnitude. Activity in this *Electome* (*electrical functional connectivity*) network is dominated by theta oscillations leading from prelimbic cortex and striatum that converge on ventral tegmental area, and by beta oscillations leading from striatum that converge on prelimbic cortex. Network activity is also synchronized with brain-wide cellular firing. Critically, this network generalizes to new groups of healthy mice, as well as a mouse line that models aberrant neural circuitry observed in brain disorders that show altered reward anticipation. Thus, our findings reveal the network-level architecture whereby the brain integrates spatially distributed activity across timescales to signal reward anticipation.

## Introduction

Anticipation of an upcoming stimulus such as a potential reward influences subsequent perception, behavior, and memory[1–3]. Human imaging studies have shown activity associated with anticipated reward in the ventral tegmental area (VTA) and nucleus accumbens (NAc), areas comprising the mesolimbic dopamine circuit canonically associated with reward response[4–6]. This anticipatory activity has also been observed across a broader set of brain regions including prefrontal cortex[7–9], hippocampus[4, 8, 10], striatum[11, 12], and thalamus[12, 13]. Pre-reward neural activation is potentiated in anticipation of greater potential rewards across several of these regions[5, 7, 14], a finding recapitulated by *in vivo* studies in animal models using electrical and neurochemical recordings of VTA and NAc[15–17]. These latter experiments, which allow for faster measurements than those possible using human imaging technology, have also revealed a ramping temporal profile of the anticipatory signals (i.e., neural signals were positively correlated with the progress of animals towards obtaining the reward)[15, 17-23].

Because a similar ramping profile in cellular action potentials occurs across primary sensory regions during periods linked to stimulus or reward anticipation in rodents[24, 25], we hypothesized that a brain-wide *Electome* network[26] coordinated across multiple timescales may broadly organize spatially distributed cellular activity to encode reward anticipation. To address this question, we implanted seven healthy inbred mice with multi-wire recording electrodes at several sites across regions that putatively signal anticipation including the prelimbic cortex (PrL, an anatomical subdivision of medial prefrontal cortex), dorsal medial striatum (DMS), medial dorsal thalamus (mdThal), dorsal hippocampus (dHipp), NAc, and VTA. We then repeatedly subjected animals to a sample-to-match T-maze assay that was modeled after a previously validated behavioral paradigm in which reward anticipation was operationalized based on the progress of rodents towards obtaining a reward[15].

We used an unsupervised machine learning analysis approach on local field potential activity acquired from the six brain sites to learn the naturally occurring organization of neural activity across multiple timescales and brain regions. We discovered multiple *Electome* networks; nevertheless, only one of these *Electome* networks signaled anticipation. This network showed ramping activity as animals approached a potential reward and, in alignment with psychometric studies in humans and rodents, the network signaled potential reward magnitude within-subject. Network activity was dominated by theta oscillations leading from prelimbic cortex and dorsal medial striatum and beta oscillations leading from striatum (dorsal medial striatum and nucleus accumbens) that converged on VTA and prelimbic cortex, respectively, and it was synchronized with brain-wide cellular firing.

Critically, the anticipation network generalized to a separate cohort of mice performing a delayed version of the sample-to-match test. Network activity in these new mice was impacted by behavioral manipulations that modulated their reward expectancy, again within subject. Finally, we found that this network also generalizes between groups of mice to signal the enhanced reward expectancy observed in a mouse model of a neuropsychiatric disease state. Taken together, these findings reveal a behaviorally-relevant brain-wide *Electome* network that is descriptive (explains feature organization), predictive (responds in an expected manner to external perturbations), generalizable (extrapolates across subjects), and convergent (extrapolates across multiple biological contexts)[27]. Thus, our findings achieve the lofty criteria for explainable artificial intelligence[27], XAI, yielding a new network-level neural mechanism whereby the brain coordinates distributed neural activity to encode reward anticipation.

## Results

In order to record neural activity during reward anticipation, we designed a fully automated sample-to-match T-Maze assay that was modeled after behavioral tasks previously used to probe the neural mechanisms underlying reward anticipation in mice[15]. The maze consisted of a stem and two arms, each of which was equipped with a nose-poke hole that could detect the presence of a mouse’s nose and dispense a liquid sucrose reward (Fig. 1a). To initiate a trial, a mouse activated a nose poke detector in the stem. After a small sucrose reward (5µL, 10% sucrose) was delivered directly into the poke detector port, a set of LED lights was turned on: LED lights on the right-hand side of the stem indicated that the mouse should proceed to the right arm of the maze to receive a reward, while LED lights on the left of the stem indicate that the mouse should proceed to the left arm. The gate then opened, allowing the mouse to run down the stem into one of the arms, where it could nose-poke at the end. A large liquid sucrose reward (15µL, 10% sucrose) was delivered into the nose poke port only if the mouse chose the arm that matched the LED light cue. Mice performed this task daily for 30 minutes or 100 trials, whichever occurred first. Daily neural recording began at P80 and continued for 35 days (Fig 1b). We then analyzed neurophysiological activity during reward anticipation, operationalized as the interval between cue presentation and reward delivery, in addition to the interval during which the mouse returned to the stem, using camera tracking to record the mouse’s goal progress (Fig. 1c, left).

**Figure 1.**
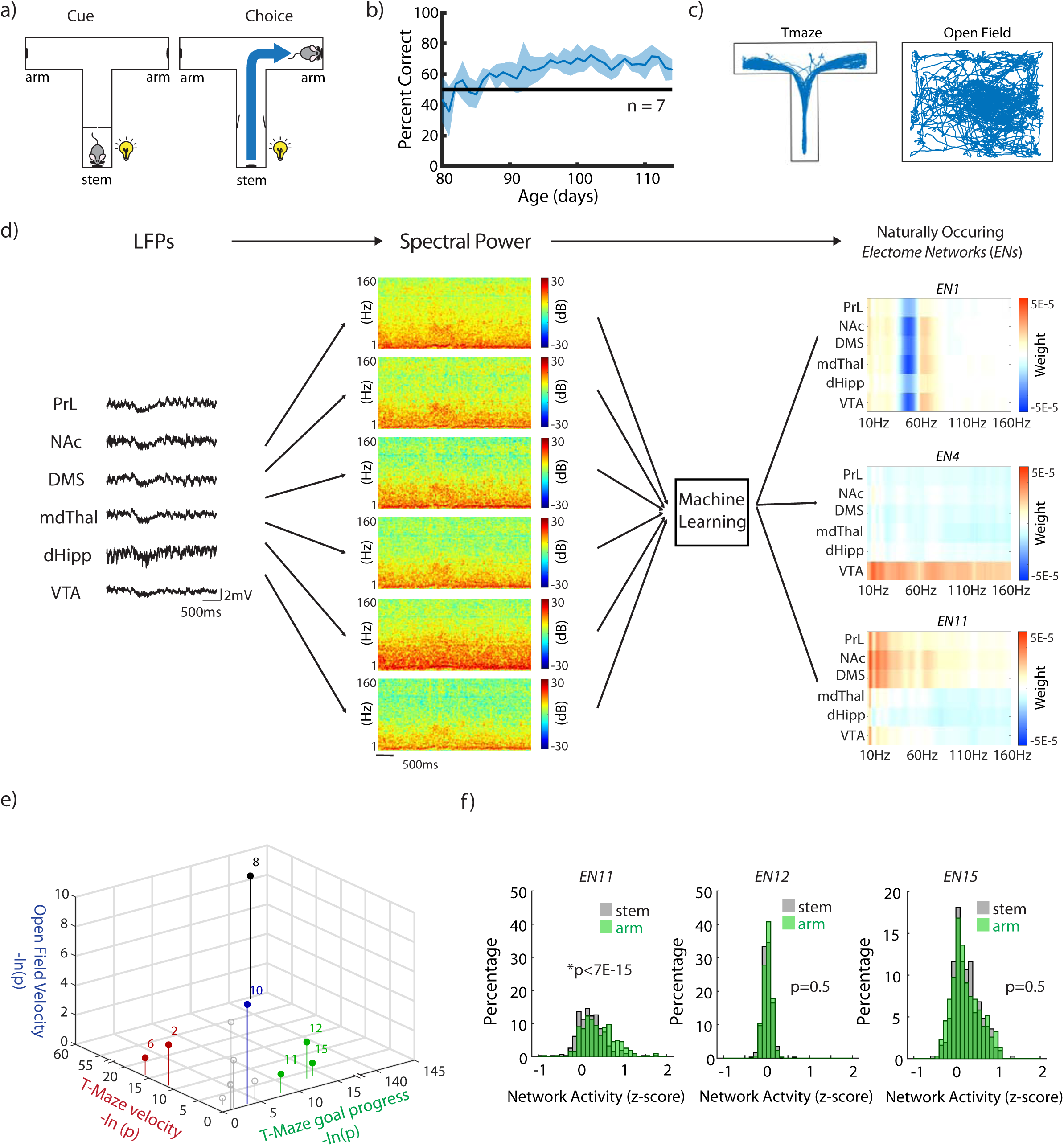
A naturally occurring electrical network that signals anticipation. **a)** Schematic of cued T-maze sample-to-match task. **b)** Task accuracy over the course of the experiment (n=7 mice). Data was sampled P80-P115. **c)** Example position-tracking from a single test day in the T-maze and open field tasks. **d)** Power spectrograms were calculated from LFPs recorded from 6 brain regions. ICA of these power spectrograms yielded 15 components, i.e., *Electome* networks (*EN*). **e)** *EN11*, 12, and 15, correlated significantly with goal progress in the T-maze without correlating with velocity in the T-maze or open field. Axes show −ln(p-value), with colored symbols representing statistical significance along the axis of the matching color. **f)** Only *EN11* showed arm>stem activity just before the goal (p<7×10^−15^ using mixed effects general linear model, testing for effect of run direction with random intercept for mouse). Data shown as mean±95%CI.

We uncovered 15 naturally occurring *Electome* networks, composed of coordinated multi-region fast activity patterns observed while *Arpc3^f/f^*:Cre-negative mice[28] (WT mice) explored the sample-to-match assay. We achieved this using Independent Component Analysis (ICA)[29-32], a well-established method of unsupervised machine learning that discovers features in large data sets (here, millisecond-resolved neural electrophysiological data) that change together over time and parses them into multiple components (i.e., *Electome* networks) (Fig. 1d). Here, we found fifteen components was the number that best-balanced complexity (i.e., prioritizing explaining more variance) with parsimony (i.e., prioritizing choosing fewer components; see Supplemental Fig. S2a-b). Thus, in contrast to prior studies that probed the role of individual brain regions or circuits within a putative anticipation network[5, 15, 17-21], this approach allowed us to interrogate many circuit elements concurrently and to discover how they are integrated across space and multiple timescales from hundreds of milliseconds to seconds. Electrophysiological signals were recorded from regions comprising multiple cell types and projections distributions; thus, we chose ICA because this machine learning model allowed each region or circuit to contribute to multiple networks, as quantified by their learned feature weight within each network/component. Additionally, each network using ICA exhibits a time-evolving activity score, which quantifies its activation strength at each interval within the observed time series (see Supplemental Fig. 2c). Critically, networks were learned independently of behavior. Since this machine learning approach is unsupervised and does not include any behavioral information, the model does not ‘encourage’ any of the learned networks to be relevant to the behavioral/cognitive process of interest (i.e. goal progress/reward anticipation).

The spatiotemporal dynamics of only one of the fifteen naturally occurring *Electome* Networks, *Electome* Network 11 (*EN11*), reflect a fundamental mechanism whereby the brain integrates spatially separated brain circuits across multiple timescales to signal anticipation. We established this finding using a clearly defined set of criteria. First, we determined whether any of our measured *Electome* network activity scores correlated with goal progress. Since velocity of mice in our sample-to-match assay was also correlated with their goal progress (Supplemental Fig. S3), we disambiguated signaling of reward anticipation from velocity by modeling each network’s activity as a linear combination of each animal’s instantaneous velocity and its progress towards obtaining a reward. Second, we further verified that the activity scores for our *ENs* did not simply reflect velocity by also subjecting the same mice to neural recordings in a novel open field, an assay which elicits exploratory behavior in the absence of a targeted external reward (Fig. 1c, right). With this approach, we found that three of our fifteen *ENs*, 11, 12, and 15, signaled goal progress in the sample-to-match assay, but not velocity in either setting (Fig. 1e). Lastly, since previous studies have established that anticipatory-brain activity is modulated by the size of the expected reward[5, 7, 15], we also tested whether any of these three networks exhibited similar psychometric responses within subject. In our task, animals ran stem→arm, responding with a nose poke for a potential large sucrose reward (15µL), and then arm→stem to return, responding with a nose poke for a small sucrose reward (5µL). We compared the strength of the three *Electome* networks immediately prior to the reward poke (i.e., 90-95% of goal progress) in the stem pokes vs. the arm pokes and found that only *EN11* showed activity that was higher in the stem→arm runs (Fig. 1f). Together, these observations advanced *EN11 a*s a putative neurophysiological correlate of anticipation: it’s signaling reflects potential reward magnitude and goal progress independent of velocity (Fig. 2a-b).

**Figure 2.**
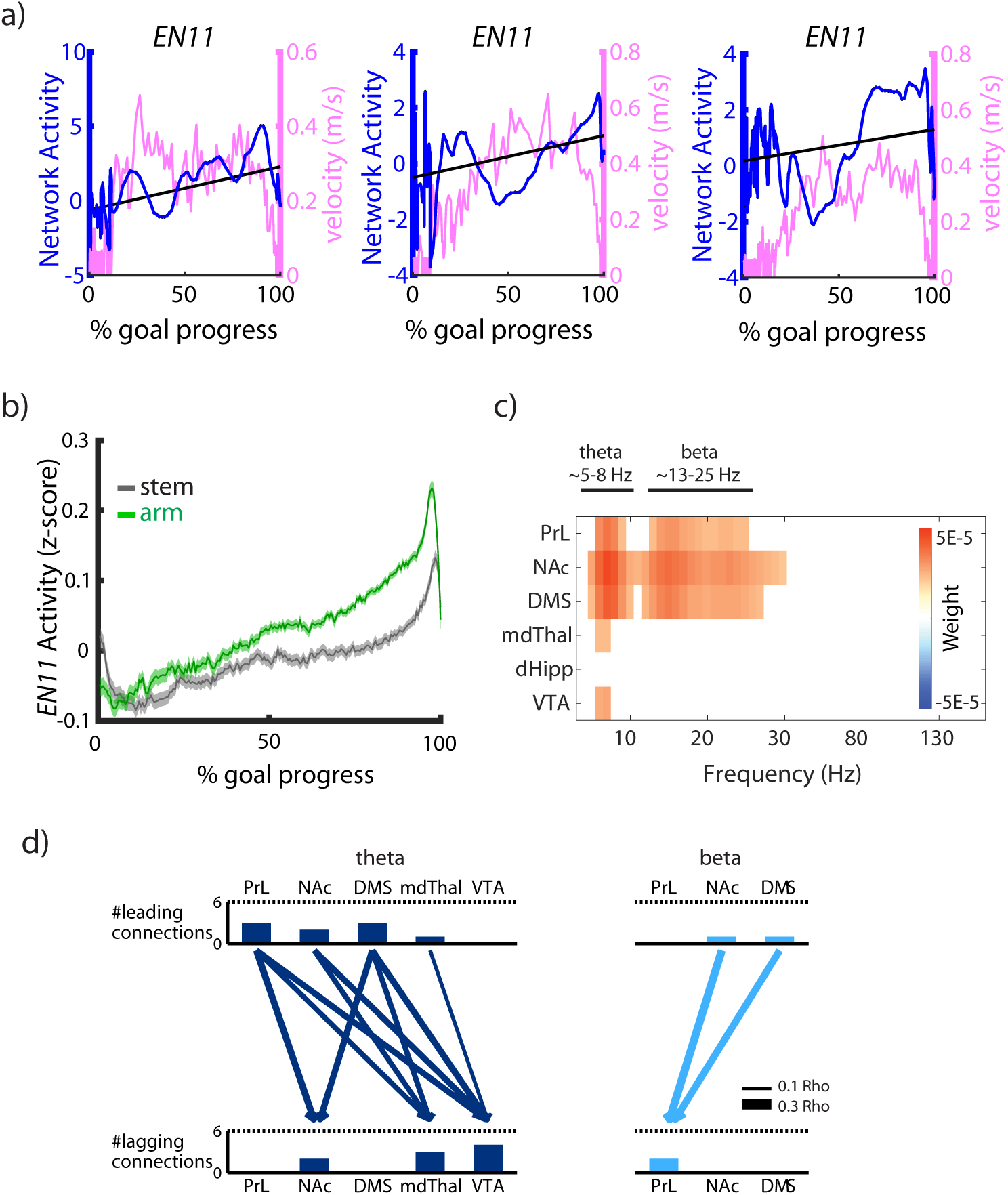
Spatiotemporal architecture of Electome Network 11. **a)** Examples of *EN11* activity (blue, axes scaled to 10^3^) and velocity (pink) for stem→arm runs on three randomly-chosen individual trials. **b)** Task specific *EN11* activity averaged over all trials. Shaded region represents 95% confidence interval across trials and mice. Notice ramping during arm→stem (grey) and stem→arm(green) runs. **c)** Thresholding at 2 standard deviations shows that *EN11* is comprised mainly of theta power in PrL, NAc, DMS, mdThal, and VTA, ad beta power in PrL, NAc, and DMS. The frequency scale has been compressed for the 30-160Hz range for visualization purposes. **d)** Directionality analysis of *EN11.* Arrows represent direction of information flow, and bar graphs above and below summarize the number of arrows leading from an area and converging on an area, respectively. In the theta band, PrL and DMS are leading areas of signal flow, while VTA is the dominant sink. In the beta band, PrL is the sink for signal from NAc and DMS. Pairwise coherence of each of these connections also significantly correlates with network activity, the strength of which is represented by arrow weight (Pearson Rho). Data shown as mean±95%CI.

The organization of the activity across the brain regions that composed *EN11* exhibited spatiotemporal directionality that converged on VTA. By thresholding the unmixing matrix (i.e., the contribution weights) at least two standard deviations in magnitude (Fig. 2c), we found that *EN11* was composed of activity predominately organized by frequency bands rather than individual regions in isolation (compare *EN11* to *EN4*, for which the latter network is largely represented by VTA activity, see Fig 1d, right): theta oscillations (~5-8Hz) in prelimbic cortex, nucleus accumbens, dorsal medial striatum, medial dorsal thalamus, and VTA, and beta oscillations (~13-25Hz) in prelimbic cortex, nucleus accumbens, and dorsal medial striatum (Fig. 2c). When we performed directionality analysis on the spatiotemporal features that composed this putative anticipation network, we discovered that the synchronous beta oscillations led from basal ganglia (dorsal medial striatum and nucleus accumbens) to prelimbic cortex. The synchronous theta oscillations in the network led from prelimbic cortex and dorsal medial striatum, relayed through nucleus accumbens and medial dorsal thalamus, and converged in VTA (Fig. 2d), providing support for prior observations that suggest anticipation may be regulated by the PFC[7, 9]. To test whether each of these circuit elements was indeed part of the broader network, we compared EN11 activity with pairwise coherence, a correlate of circuit activation quantified by synchronous millisecond timescale fluctuations in neural activity. Indeed, for all pairs of regions within *EN11*, coherence within the theta and beta frequency bands was significantly correlated with *EN11* activity at the supra-milliseconds timescale (p<0.001 for all comparisons of pairwise coherence and *EN11* activity using a mixed effects general linear model, testing for the group mean correlation coefficient > 0, with random intercept for mouse; Fig. 2d). Nevertheless, activity in the independent regions/frequency bands (i.e., theta or beta frequency spectral power) did not correlate with goal progress on their own (Supplemental Fig. S2d), providing evidence that anticipatory signaling by *EN11* represents an emergent signal that arises from the coordination of many individual circuit elements, rather than from any specific circuit element in isolation. Thus, the spatiotemporal dynamics represented by *EN11* reflect a fundamental mechanism whereby the brain integrates spatially separated brain circuits across multiple times scales to signal reward anticipation.

*EN11* activity also reflects the spatiotemporal dynamic mechanisms that coordinate cellular activity across brain regions and timescales to encode reward anticipation. We identified cellular activity across multiple brain regions that exhibited ramping as animal progressed towards the reward (Fig. 3a-b, Supplemental Fig S4). To probe the relationship between *EN11* activity and network-wide cellular firing, we measured and compared cellular firing and *EN11* activity during all stem→arm runs. Here, we quantified both measures within equally segmented bins, based on the percent goal progress from 5%-95%, for each stem→arm run. After directly correlating cellular firing and *EN11* network activity across all the bins for a given session, we created a null distribution by shuffling our cellular activity bins across trials, while maintaining the relationship between cell firing and goal progress. Since our results and prior studies had shown ramping anticipatory cell firing in multiple regions[17, 18, 20, 22, 24, 25], this approach allowed us to directly probe whether cellular firing signaled *EN11* activity in a manner that could not simply be explained by the observation that both measures signal goal progress independently. Using this strategy, we discovered cells in every brain region that correlated with *EN11* activity in WT mice (Fig. 3c).

**Figure 3.**
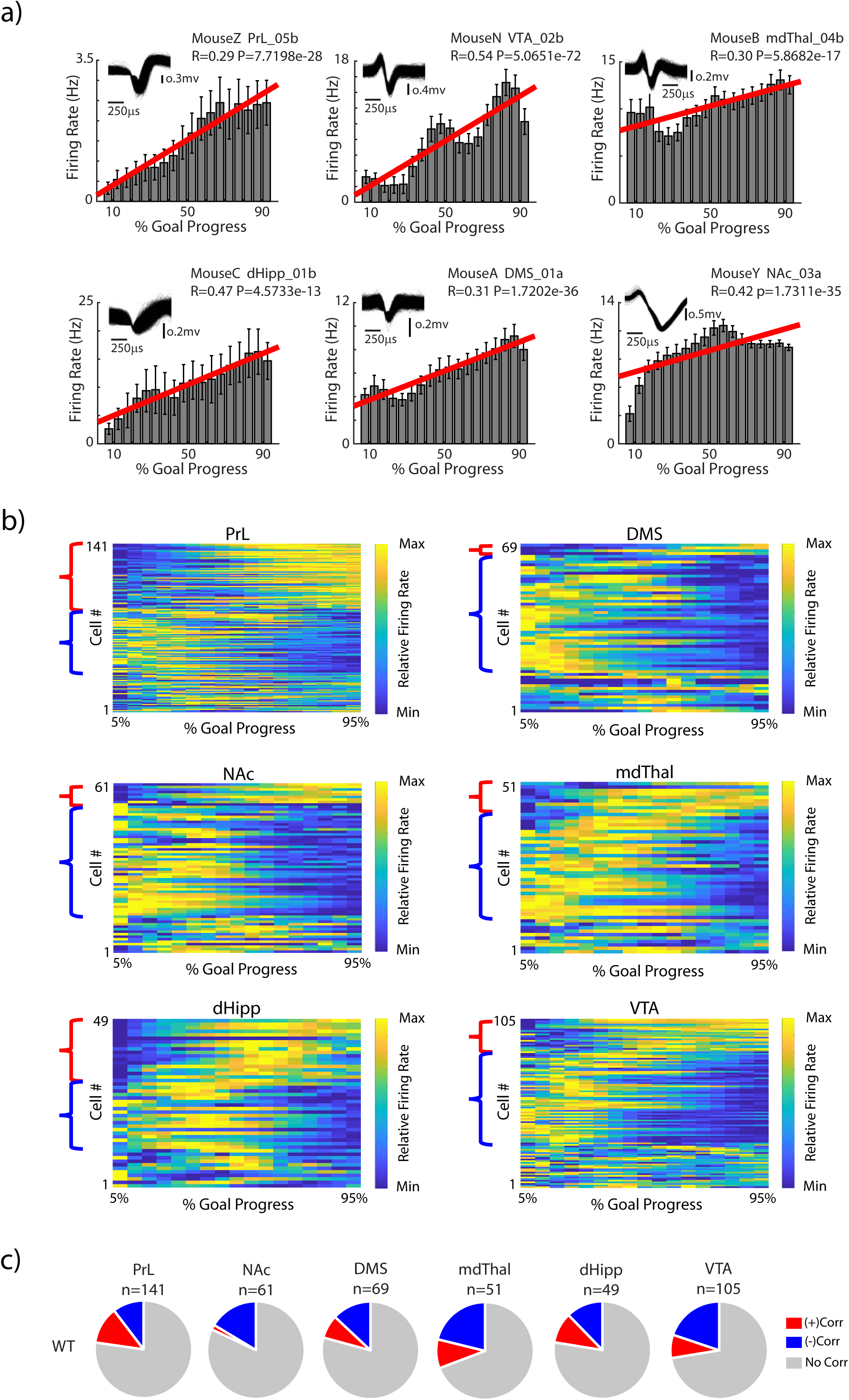
Electome Network 11 activity reflects network-wide cellular activity. **a)** Histograms depicting representative examples of cells from each area that exhibit mean firing rates positively correlated with goal progress for stem→arm runs (i.e., ramping). Regression lines reflecting the significant correlations are shown in red. Firing rates were averaged across all trials within a single session. Insets show spike waveforms. **b)** Relative mean firing rate of cells recorded from each brain area versus goal progress. Firing rates were averaged across all trials within a single session. Red brackets highlight cells which showed firing rates positively correlated with goal progress (i.e., ramping), and blue brackets highlight cells which showed firing rates that were negatively correlated with goal progress (i.e., sledding). **c)** Proportion of cells positively (+, red) and negatively (-, blue) correlated with network activity in WT mice.

Next, we verified that *EN11* indeed signaled anticipation using a gold-standard machine learning validation strategy[27]. Specifically, we tested whether the *EN11* network learned from our original group of WT mice signaled anticipatory behavior in a separate group of normal healthy mice. LFP spatiotemporal dynamics acquired from a new group of C57BL/6J mice (C57) were projected into our *Electome model* using the ICA component loadings learned from our original analysis. Here, however, the new group of implanted mice were subjected to a modified version of our sample-to-match task that included delays ranging from 0.5s to 8s after the cue presentation in the stem, during which animals were unable to advance towards the arms (delayed sample-to-match) (Fig. 4a). In this task, after cue presentation in the stem, mice performed a nose poke under the LED cue, which turned off the cue and began the delay (Fig. 4a, top right). Following the delay, the gate opened, and mice could proceed to make their response in the arm (Fig. 4a, bottom). Mice underwent the same training as mice in the original task, with the addition of training stages to ensure they could perform the task with the delay (see Supplemental Methods). During the subsequent task performance, the accuracy of mice in the delayed sample-to-match task (i.e., proportion of correct trials) decreased with increasing time delays (Fig. 4b). When we projected LFP activity acquired from this validation set of mice during this modified sample-to-match assay into the *Electome* Network model (ICA space) learned from our original data set, we found that the model explained at least as much variance in the validation data set as it had in the original data set (variance explained: original dataset, 20.1±0.6; delayed sample-to-match data set, 36.5±2.0; mean±sem; p > 0.99, one-sided unpaired t-test). *EN11* ramped towards goal progress in the C57 mice and ramped more for the stem→arm runs than for the arm→stem runs for the 0.5s time delay, providing clear validation for our initial observations (p = 0.047, using a mixed effects general linear model of EN11 value averaged over 90-95% goal progress as a function of 0.5s-delay arm→stem vs. stem→arm runs. with random intercept for mouse; see Fig. 4c-d).

**Figure 4.**
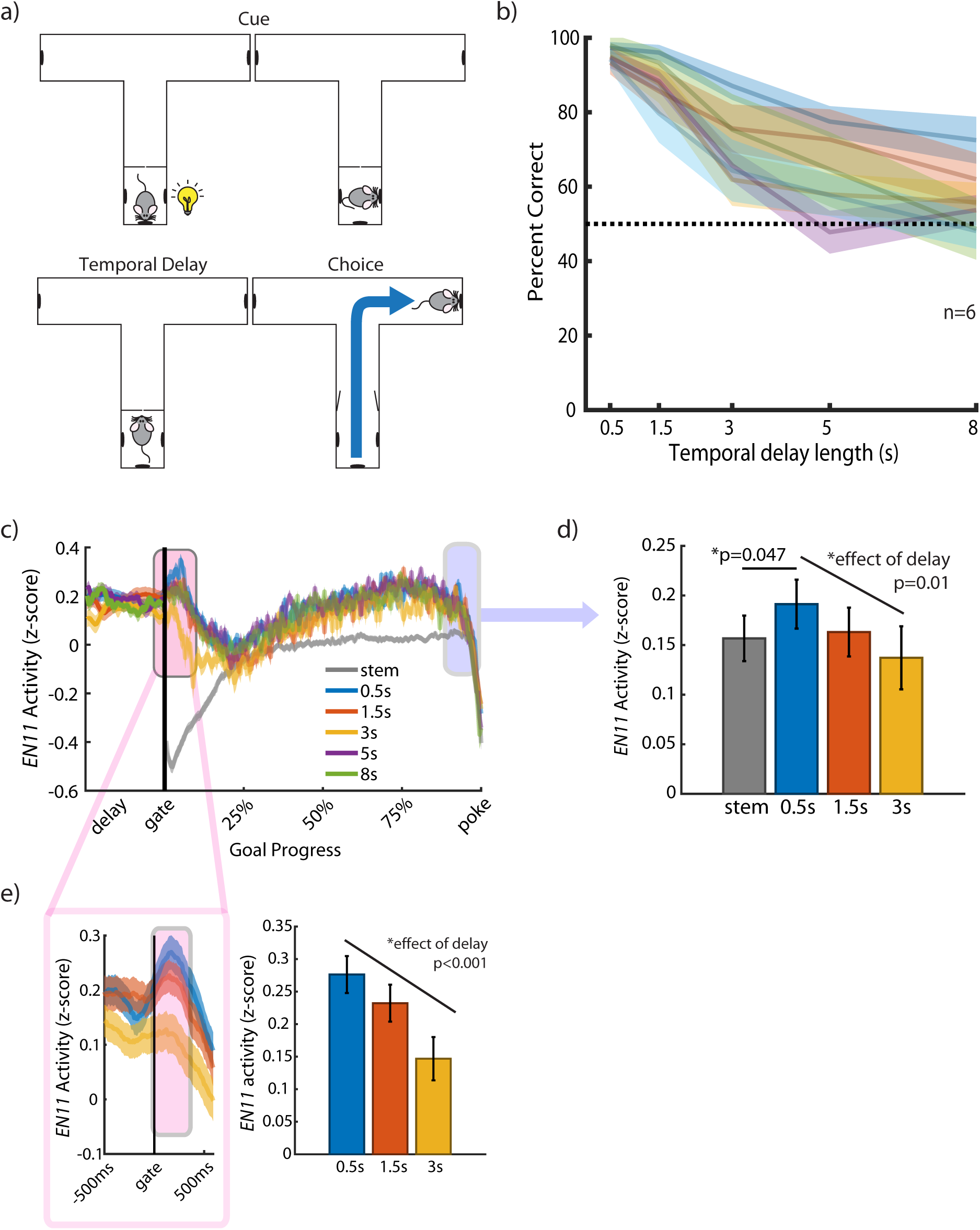
Electome Network 11 activity signals reward anticipation in a new cohort of mice. **a)** Schematic of delayed sample-to-match task. **b)** Task performance accuracy across mice (n = 6 mice). Each line reflects a separate mouse, with shaded region corresponding to 95% confidence interval. Task performance was above chance for 0.5, 1.5, and 3s temporal delays. **c)** *EN11* activity during the task, averaged across mice and trials. The shaded region represents the 95% confidence interval. Mean stem→arm activity at 90-95% goal progress in the 0.5s delay, the closest to a direct replication of the original sample-to-match task, is greater than mean stem→arm activity at 90-95% goal progress (note that arm→stem activity is plotted such that the ‘gate’ corresponds with the prior nose poke in the arm) **d)** There is a negative correlation between *EN11* activity just before goal and delay length, and for **e)** for *EN11* activity just as the mouse leaves the gate.

Next, we asked whether *EN11* activity was directly modulated by reward expectation in the out-of-sample validation testing C57 animals. We exploited our observations at various delays in the delayed sample-to-match task to probe this relationship. We predicted that the mice, as a result of extensive training, would have a subjective estimate of reward expectation that was related to the difficulty of the trial, i.e., the length of the delay prior to the gate opening, for trials they could perform above chance (i.e., 0.5s, 1.5s, 3s, Fig. 4b). We thus hypothesized that this subjective estimate would be reflected by *EN11* activity and specifically that *EN11* activity would be negatively correlated with the length of the delay interval across these trials. Critically, each of these trials began and ended in the same location; thus, this strategy also allowed us to disambiguate the relationship between *EN11* activity and reward expectancy from additional confounding behavioral variables including the location and running path of mice, unlike for our stem→arm vs. arm→stem run comparisons in the initial WT animals. When we compared *EN11* activity in these new mice immediately before the reward poke (90-95% goal progress) across the 0.5, 1.5, and 3s delay trials, we found that *EN11* activity at 90-95% goal progress decreased as the time delays increased (and task accuracy decreased), supporting our hypothesis (p=0.01 using a mixed effects general linear model of mean *EN11* activity at 90-95% goal progress as a function of delay, with a random intercept for mouse; Fig. 4c-d). Though uncertainty has also been shown to increase reward anticipatory activity[17], these findings confirmed that the increased *EN11* activity observed in the initial sample-to-match task at the end of stem→arm runs (~65% rewarded) vs. the stem→arm runs (100% rewarded) was not solely due to uncertainty. Specifically, higher EN11 activity was observed at for the stem→arm vs. arm→stem runs at 0.5s delays where ~95% of trials were rewarded, and *EN11* activity was lower at delays with increased uncertainty (0.5s vs. 1.5s and 3s delays). Interestingly, we also observed a peak in network activity immediately after the gate opened that showed a negative correlation with delay length for the trials the mice performed above chance (p<0.001 using a mixed effects general linear model of mean *EN11* activity at 0-5% goal progress as a function of delay, with a random intercept for mouse; Fig. 4e). These results demonstrated that the *EN11* activity signaled reward expectancy within subjects at task epochs important for advancement toward a goal (i.e. mice running past the open gate and progressing towards the stem). Furthermore, these results also demonstrated that *EN11* activity did not simply reflect a motor action plan driven by prelimbic cortex to achieve a reward[33], given that the same motor sequence had the potential to yield rewards for all of the delays.

Altered anticipation has been described in disorders such as bipolar mania, schizoaffective disorder, and Parkinson’s disease in the setting of dopamine replacement pharmacotherapy. These distinct disease states are marked by both impulsivity and increased risk taking, which is thought to be linked to striatal hyperdopaminergia. Thus, we hypothesized that enhanced striatal dopaminergia would potentiate activity in the reward anticipation network, *EN11*, providing a network-level mechanism for the altered behaviors observed in these disease states. To probe this question, we exploited a conditional knockout mutant (MU) mouse line (*Arpc3^f/f^:Camk2a-Cre; littermates of the Arpc3^f/f^*-WT)[28]. These mice have a conditional knockout of the actin-related protein 2/3 complex (Arp2/3), and they have been previously shown to exhibit striatal hyperdopaminergia[34]. Notably, this striatal dopaminergia is mediated at least in part by a cortical→VTA circuit, mirroring the gross directionality we discovered in *EN11*. Mutant mice exhibited enhanced behavioral performance in our sample-to-match assay (Fig. 5a).

**Figure 5.**
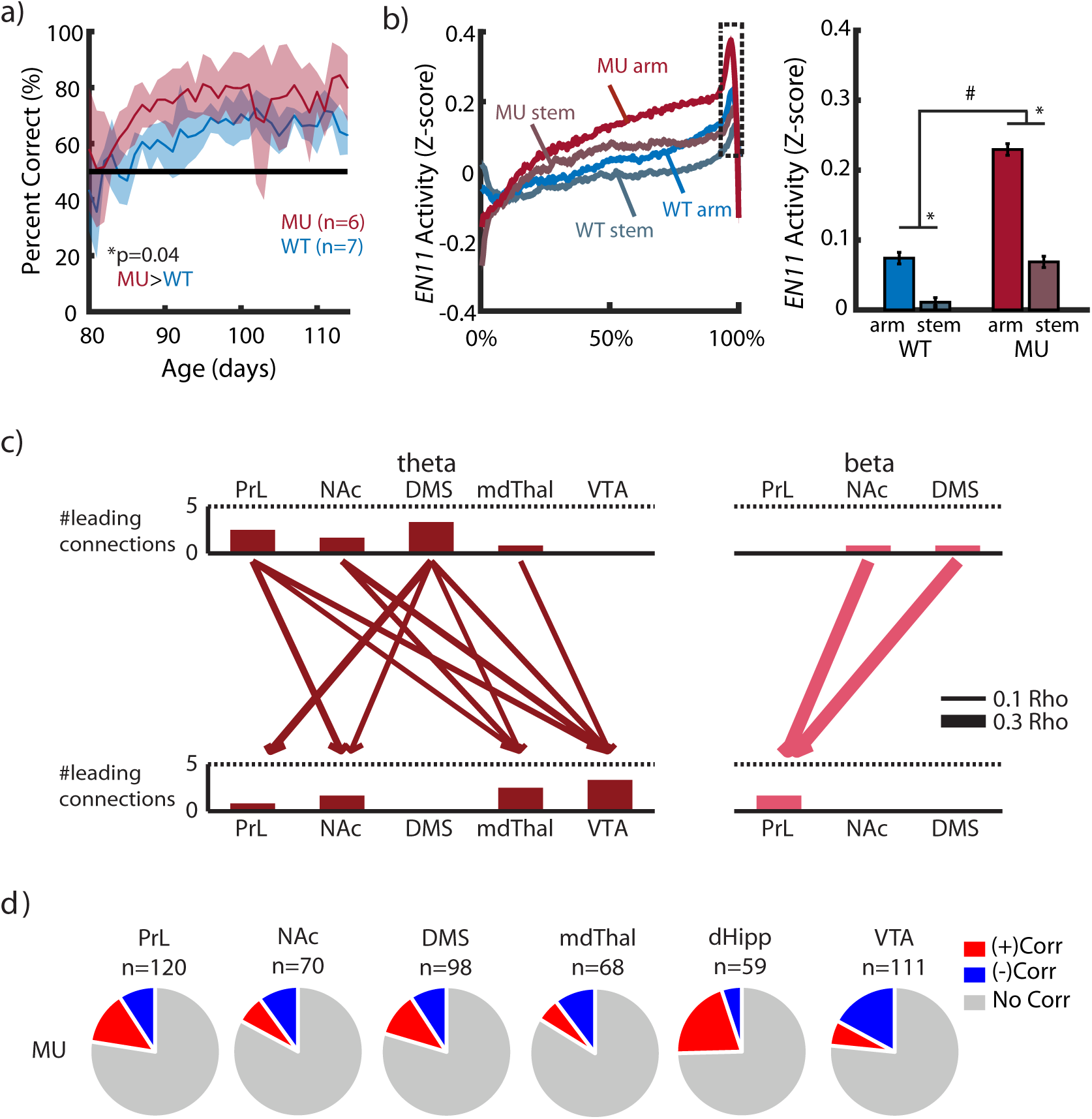
Electome Network 11 activity decodes disease states across groups of mice. **a)** Mutant (MU) mice (n = 6 mice) show increased task accuracy compared to WT mice (n=7; p=0.0003 using a mixed effects general linear model of task accuracy as a function of genotype, with a random intercept for mouse). **b)** Mutant mice show higher ramping of the network (left), both in stem→arm and arm→stem runs (#p<=0.01 and *p<0.001 for genotype and run-type effects, respectively, using a mixed effects general linear model of *EN11* activity at 90-95% goal progress, with a random intercept for mouse; right). **c)** Directionality analysis reveals the same network structure as the WT (compare to Fig. 2b), with the addition of a directional DMS-to-PrL theta connection. As in WT, pairwise coherence of each of these connections also significantly correlates with network activity, the strength of which is represented by arrow weight. **d)** Proportion of cells positively (+) and negatively (-) correlating with network activity in mutant mice.

When we projected LFP activity acquired from mutant mice during the initial sample-to-match assay into the *Electome* Network model (ICA space) learned from our initial recordings in WT mice, we found that the model explained at least as much variance in the mutant mice as it had in their WT counterparts (variance explained: original data set, 20.1±0.6 (SEM); mutant data set, 25.6±2.2; mean±sem; p = 0.99, one-sided unpaired t-test). Consistent with our initial out-of-sample validation approach, *EN11* activity in the mutant mice also exhibited ramping with goal progress, and stem→arm runs induced higher network ramping than arm→stem runs [p<0.001, using a mixed effects general linear model of *EN11* activity averaged over 90-95% goal progress as a function of run direction (arm vs stem) and random intercept for mouse; Fig. 5b]. We found directionality in the mutant network structure that was very similar to that of WT, with the addition of a directional DMS→PrL connection (Fig. 5c), and we identified cells in each brain area that signaled *EN11* activity in mutant mice (Fig. 5d). These results provided further validation that *EN11* generalized to datasets outside of the original training set, since the network signaled anticipation in the mutant mice as well. Importantly, the mutant mice exhibited significantly elevated *EN11* in both arm→stem and stem→arm runs compared to WT mice [a mixed effects general linear model of EN11 activity averaged over 90-95% goal progress as a function of genotype (mutant vs. WT) and run direction (arm vs. stem), shows both a genotype effect; Mutant>WT, p=0.01; and an run direction effect; arm>stem, p<0.001; follow up 2-sample t-tests show that Mutant>WT in arm runs, p=0.01; and in stem runs, p=0.004; Fig. 5b]. Taken together, these results showed that mice previously shown to exhibit striatal hyperdopaminergia also show enhanced activity in our putative anticipatory network. Furthermore, our findings showed that *EN11* activity could indeed be used to decode anticipation states across groups, and not only within subjects as demonstrated by our initial results in the WT and C57 mice.

## Discussion

Here, we set out to discover the network-level mechanisms whereby the brain encodes reward anticipation. By querying the organization of electrical neural signal across multiple timescales and brain regions using machine learning, we discovered a network that signaled reward anticipation, *EN11*, which was led by theta oscillatory activity in prelimbic cortex and dorsal medial striatum converging in VTA, and by beta oscillatory activity in dorsal medial striatum and nucleus accumbens converging in prelimbic cortex. This network generalizes to new animals, and psychometric manipulations that impacts reward-related behavior. Thus, our findings provide evidence for anticipation as a brain-wide phenomenon, and they demonstrate that *EN11* reflects a fundamental mechanism whereby the brain integrates activity across timescales and space to signal anticipation.

Our results also show that *EN11* activity can be used to directly decode distinct anticipation states across groups of mice. We have previously promoted our Arp2/3 conditional knockout mutant mouse line as a model of schizophrenia due to their neurophysiological and behavioral abnormalities including altered spine pruning, striatal hyperdopaminergia, hyperlocomotion, increased behavioral repetition, disrupted sensorimotor gating, and decreased rapid alternation in a classic working memory assay[28]. However, here we find that these same mice exhibit enhanced behavioral function on a sample-to-match assay and increased activity in their *EN11* anticipation network, compared to their healthy littermate controls. Indeed, since multiple disease states such as schizophrenia, schizoaffective disorder, bipolar mania, and Parkinson’s disease in the setting of dopamine replacement pharmacotherapy exhibit a profile of alterations including the aforementioned behaviors, these findings suggest that our *Electome* mapping approach may provide clearer insights regarding the phenotypes and mechanisms of neuropathology states in animal models than behavioral observations in isolation. The findings in these mutant mice, combined with the VTA-convergent signaling architecture revealed here by multisite neuroelectrophysiological recording, suggest involvement of canonical VTA dopaminergic reward circuitry. Future studies may reveal mechanisms whereby *EN11* and VTA dopaminergic signaling interact to sustain motivated behavior.

## Author Contributions

Conceptualization, MTV, SS, RAA, and KD; Methodology, MTV, LKD, GET, NMG, CBl, DNH, IHK, SDM, and KD; Formal Analysis, MTV, NMG, and KD; Investigation, MTV, LKD, GET, Cbu, MV, CBl, JW, NN, EA, and KD; Resources, IHK; SS; KD; Writing – Original Draft, MTV, RAA, and KD; Writing – Review & Editing, MTV, CBu, JW, NMG, EA, SDM, RAA, and KD, Visualization, MTV, GET, and KD; Supervision, KD; Project Administration, KD; and Funding Acquisition, MTV, RAA, KD; (see Supplemental Table S1 for experiment specific contributions).

## Declaration of Interests

The authors have no competing financial interests.

## Acknowledgements

We would like to thank S. Lisberger, M.G. Caron, K. Heller, A. West, R. Hultman, and D. Carlson, J. Schaich Borg for comments on this work; A. Talbot and B. Katz for technical support. This work was supported by NSF grant DGF1106401 and the Duke University Katherine Goodman Stern Dissertation Fellowship to MTV; NSF IIS-1451017 –BRAIN EAGER grant to RAA, KD and Katherine Heller; and a W.M. Keck Foundation grant to KD and Fan Wang. A special thanks to Freeman Hrabowski, Robert and Jane Meyerhoff, and the Meyerhoff Scholarship Program.

