## Supplemental Materials for "Brain-wide electrical spatiotemporal dynamics encode reward anticipation"

### Supplemental Table

| | |
|---|---|
| Mai-Anh T. Vu | Jointly conceived all experiments. Jointly designed and built behavioral apparatus used for all experiments and jointly developed methodology for behavioral tasks. Jointly collected neurophysiological, behavioral, and video data for open field, sample-to-match, and delayed sample-to-match experiments. Jointly processed video data for all experiments. Jointly analyzed data for all experiments. Wrote and revised the manuscript. |
| Lisa K. David | Jointly developed training methodology and collected data for sample-to-match and delayed sample-to-match experiments. Jointly developed behavioral apparatus. |
| Gwenaëlle E. Thomas | Jointly collected data for sample-to-match experiment. Jointly built behavioral apparatus used for all experiments. Jointly performed histology (slicing, mounting, imaging) for all experiments. |
| Caley Burrus | Jointly sorted cells, collected data, and processed video data for sample to match experiments. Jointly collected data for sample-to-match experiments. Wrote the manuscript. |
| Meghana Vagwala | Jointly sorted cells, processed video, and performed histology for the sample-to-match experiment. |
| Neil M. Gallagher | Jointly developed behavioral apparatus. Assisted with machine learning analysis of neurophysiological data and performed histology for delayed sample-to-match experiment. Revised the manuscript. |
| Cameron Blount | Built electrodes and jointly collected data for delayed sample-to-match experiment. Jointly performed histology (slicing, mounting) for all experiments. |
| Joyce Wang | Jointly collected data for sample-to-match and delayed sample-to-match experiments. Revised the manuscript. |
| Dalton N. Hughes | Jointly developed behavioral apparatus used for experiments. |
| Nkemdilim Ndubuizu | Jointly collected data for delayed sample-to-match experiment. Jointly performed histology (slicing, mounting) for all experiments. |
| Elise Adamson | Jointly collected data for delayed sample-to-match experiment. Revised the manuscript. |
| Il Hwan Kim | Jointly designed experiments and contributed the mutant mouse line; genotype verification. |
| Scott Soderling | Jointly conceived sample-to-match experiments. Developed and contributed mutant mouse line. |
| Stephen Mague | Designed, built, and implanted recording electrodes for sample-to-match experiments; supervised histology (slicing, mounting) for all experiments. Wrote and revised the manuscript. |
| R. Alison Adcock | Jointly conceived sample-to-match experiments. Wrote and revised the manuscript. |
| Kafui Dzirasa | Jointly conceived all experiments. Jointly designed behavioral apparatus and training methodology, for sample to match and delayed sample-to-match tests. Designed and implanted recording electrodes for delayed sample-to-match experiment. Supervised data collection and processing for all experiments. Jointly analyzed cellular data. Wrote and revised the manuscript. |

**Supplemental Table S1.** ***Detailed Author Contributions***

### Supplemental Figures

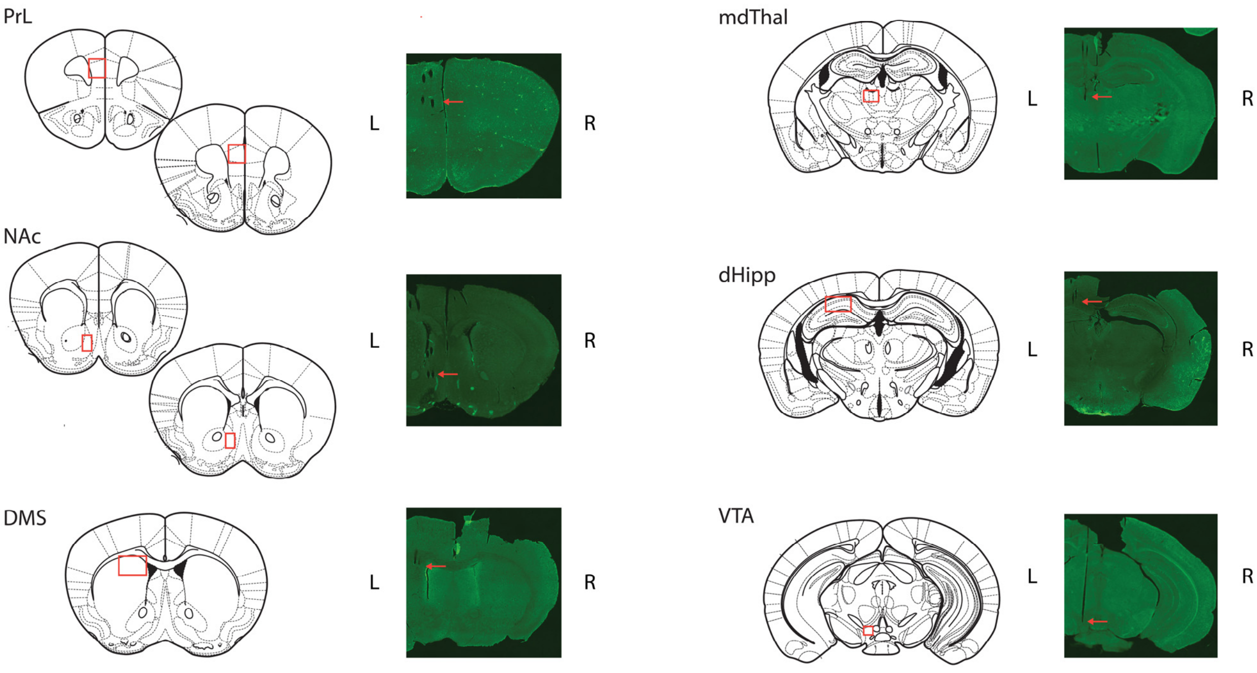


**Supplemental Figure S1.** ***Histological confirmation of electrode placements***

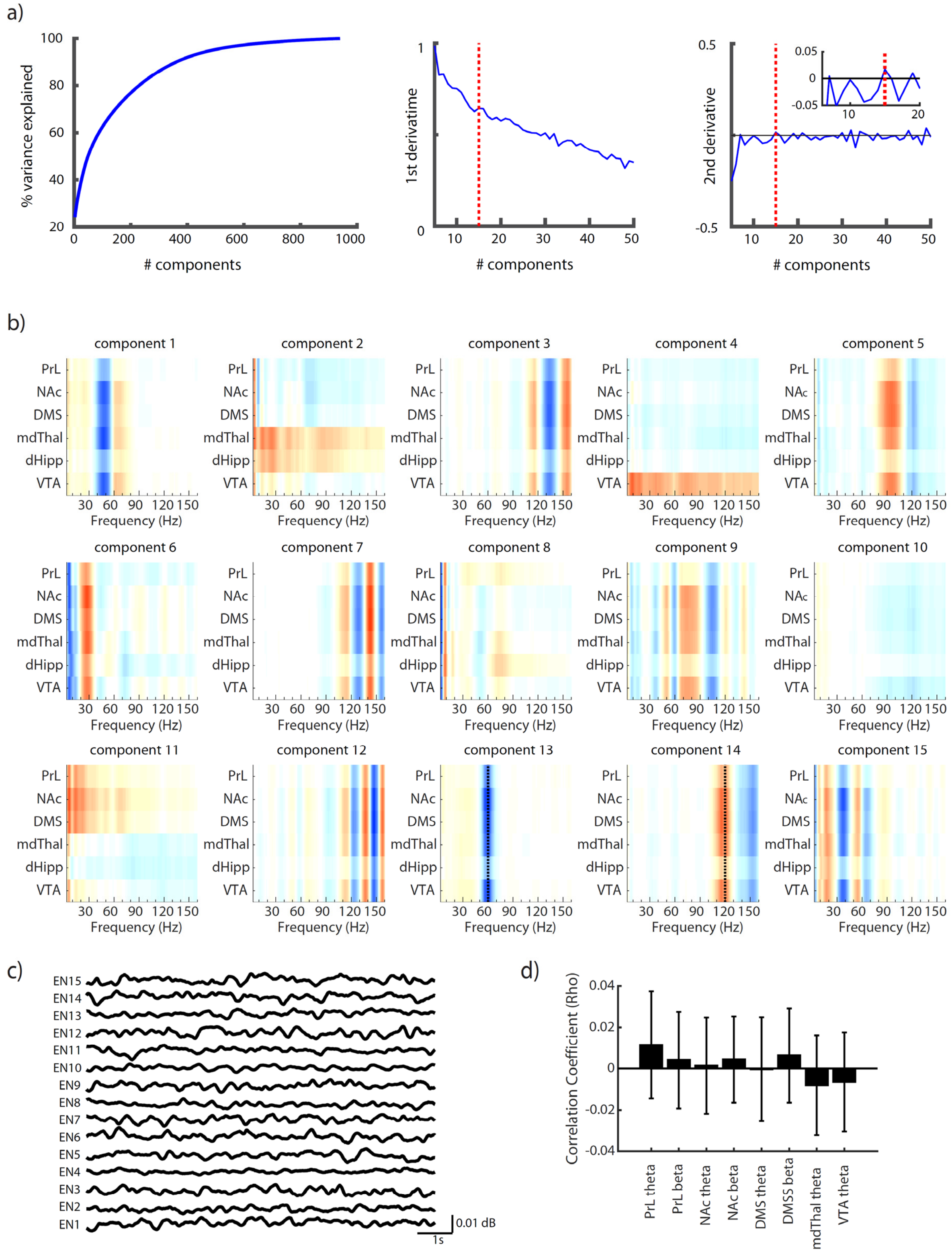


**Supplemental Figure S2.** ***Discovering naturally occurring (observed) networks using Independent Component Analysis.*** **a)** Left) The training data was modeled with varying numbers of components, from 3 to 942, the maximum number of components (in this case, each component is composed by a single 1Hz frequency band from a single region). A separate set of validation data was projected into these components, and the % variance explained calculated. This % variance explained increases as the number of components increases. The elbow method was used to balance complexity (prioritizing % variance explained) with parsimony (prioritizing a smaller number of components). The objective is to identify the number of components after which the gain in % variance explained lessens. Middle) The first derivative of the % variance explained vs. # components, representing the gain in % variance explained for each additional component included in the model. This value decreases with additional components. Right) The second derivative of the % variance explained vs. # components. The point where the 2nd derivative crossed 0, indicates a flattening out of the change in % variance gained for each additional component included in the model. Inset shows the $2^{nd}$ derivative at around 15 components. **b)** The learned ICA components. Note that several networks captured spatiotemporal dynamics related to signal artifact (i.e. 60Hz electricity signal). Rather than remove these networks from further analysis, we chose to probe whether the networks that survived our behavioral/psychometric criteria for signaling anticipation also exhibited physiologically relevant spatiotemporal dynamics (i.e., correlations within cellular firing, and network directionality). Only component 11 (Electome Network 11, *EN11*) survived our initial inclusion criteria and passed the subsequent neurophysiological tests as well. **c)** Sample activity traces for learned Electome networks. **d)** Spectral features within *EN11* failed to signal anticipation in isolation when they were directly compared to goal progress ($p>0.5$ for all comparisons using a mixed effects general linear model, testing for the group mean correlation coefficient > 0, with random intercept for mouse).

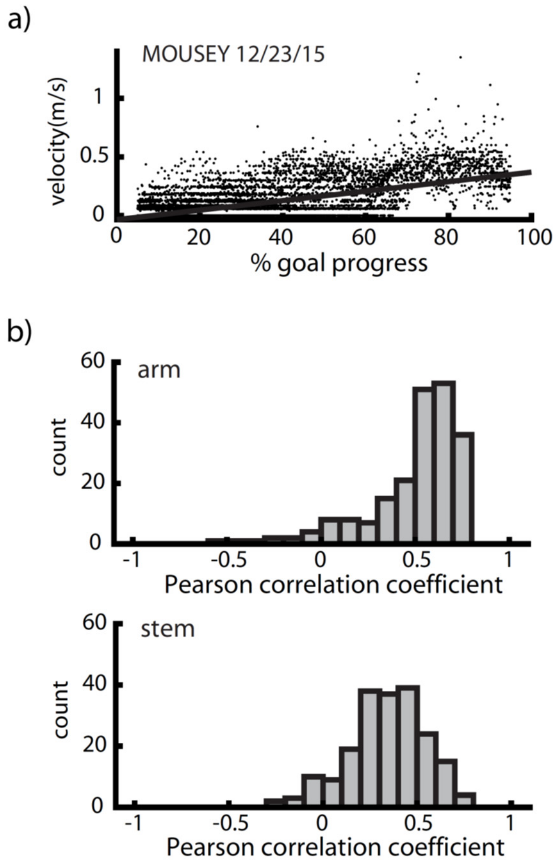


**Supplemental Figure S3.** ***Relationship between velocity and goal progress in the sample-to-match task.*** **a)** Scatter plot of velocity vs. goal progress for a single recording session from a single mouse. **b)** Distribution of correlation coefficients between velocity and goal progress in stem→arm runs, across mice and test sessions. A mixed general linear model of group mean correlation coefficient with random intercept for mouse shows that this is significantly different from 0, $p<0.001$. **c)** distribution of correlation coefficients between velocity and goal progress in arm→stem runs, across mice and test sessions. A mixed general linear model of group mean correlation coefficient with random intercept for mouse shows that this is significantly different from 0, $p<0.001$.

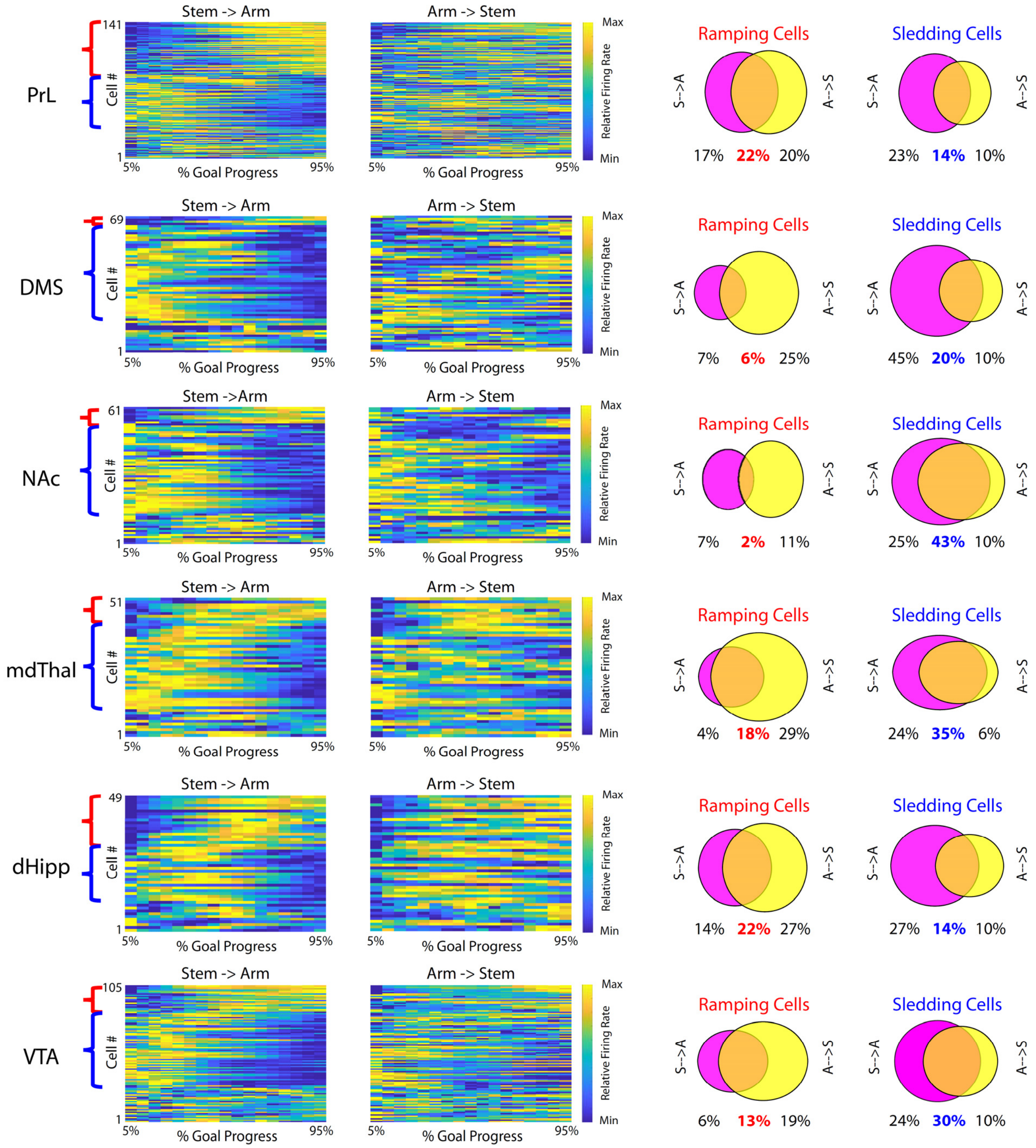


**Supplemental Figure S4. *Correlated activity between cellular firing and goal progress.*** Relative firing rates vs. goal progress for recorded cells during stem→arm runs and arm→stem runs. Cells that which showed positive correlations between their relative firing and goal progress for both run types were verified as ramping; cells that showed negative correlations between their relative firing and goal progress for both run types were verified as sledding. Cells that showed correlations under one run type both not another may reflect other task-relevant variables, such as the location of mice in the maze.

### Supplemental Methods

#### Animal Care & Use

Nineteen male mice were used in this study: six conditional knockout Arp2/3 mutant mice (*Arpc3$^{f/f}$:Camk2a-Cre*), seven Cre-negative littermates (*Arpc3$^{f/f}$*), and six wildtype (C57BL/6J) mice. The Arp2/3 mutant mice and Cre-negative littermates were generated as described previously[1], and backcrossed six generations onto a C57BL/6J background. All mice were group-housed 3-5 mice per cage in the Duke University Division of Laboratory Animal Resources facilities on a 12-hour light/dark cycle and maintained in a humidity- and temperature-controlled room with water available ad libitum. Except when food-restricted for the purpose of behavioral training and testing, mice were given ad-libitum access to food. Food-restriction did not occur until mice were adults of age at least 9 weeks (P63) and at least 2 weeks post-electrode implantation surgery. During food restriction, mice were gradually reduced to ~90% of their free-feeding body weight[2,3]. Behavioral and electrophysiological experiments were conducted during the light cycle. All studies were conducted with protocols approved by the Duke University Institutional Animal Care and Use Committees and were in accordance with the National Institutes of Health guidelines for the Care and Use of Laboratory Animals. Note that because we expected Arp2/3 mutant mice to exhibit some progressive cognitive deficits starting by P90[1], our electrode implantation, training, and testing schedules were tightly constrained by mouse age. The genotype of these experimental animals was re-confirmed after testing via tissue sequencing.

#### Electrode Implantation Surgery

The electrode implantation surgery has been described previously[4]. Mice were anesthetized with 1.5% isoflurane, placed in a stereotaxic device, and metal ground screws were secured above the cerebellum and anterior cranium. Thirty-two tungsten microwires were arranged in array bundles designed to target prelimbic cortex (PrL), nucleus accumbens (NAc), dorsal medial striatum (DMS), mediodorsal thalamus (MdThal), dorsal hippocampus (dHipp), and ventral tegmental area (VTA), all on the left. Bundles were centered on stereotaxic anterior-posterior (AP) and medial-lateral (ML) coordinates (in mm) measured from bregma, while dorsal-ventral coordinates were measured from the dura. Number of wires, wire diameter, and coordinates were: PrL: 8×35μm wires, 1.85 AP, 0.375 ML, -1.75 DV; NAc: 4×50μm wires, 1.225 AP, 0.625 ML, -4.25 DV; DMS: 8×50μm wires, 0.75 AP, 1.125 ML, -2.25 DV; mdThal: 4×35μm wires, -1.475 AP, 0.325 ML, -3 DV; dHipp: 4×35μm wires, -2.175 AP, 1.375 ML, -1.75 DV; VTA: 4×50μm wires, -3.125 AP, 0.525 ML, -4.5 DV. For the six WT mice being tested in the delayed StM task, only three wires were implanted in NAc and DMS. Implanted electrodes were anchored to round screws using dental acrylic. Recording sites were confirmed histologically at the conclusion of experiments.

#### T-Maze Apparatus

The T-maze apparatus, constructed from black plastic Legos, had approximate dimensions of 48cm wide x 35cm deep x 30cm tall, where 48cm was the arm-to-arm distance, and 35cm the stem-to-back-wall distance. The arms and stem were approximately 10cm wide. The gates in the middle of stem were approximately 12cm from the front of the stem. The maze was equipped with five nose poke holes, which detect nose-poke via IR beam breakage. The nose poke holes were located at the end of each arm (2), at the end of the stem (1), and by each gate (2). The maze had four 4-LED light columns: one on either side of the stem nose poke hole and one on each side just before the gate. Lighting was dim: 3-30 lux. Earphones on the back wall were calibrated to deliver 68dB white noise. The task (i.e., data recording, timing, lights, gates, reward, and gates) was fully automated.

**Behavioral Training & Testing**

Mice were habituated to handling and the T-maze for five days prior to electrode implantation surgery; during this time, mice were put in the maze with their cage mates, and all nose pokes into nose poke holes were rewarded with 5µL of 10% sucrose solution. Mice recovered from electrode implantation surgery for at least five days before being habituated to the maze again and commencing training, though not food-restricted until at least two weeks post-surgery. Mice began light behavioral training five days after surgery, and after two weeks, were habituated to being plugged into an acrylic “dummy plug”, matching the size and weight of the recording set up, which the mice wore during training. This continued until behavioral testing so that mice were fully accustomed by the time neurophysiological recording began.

**Sample-to-Match T-Maze Task**

To start each trial, the mouse poked the stem to start each trial, getting rewarded 5µL of 10% sucrose solution. At this time, the gate closed for two seconds, while either right or left LED light cues turned on. Once the gate opened, the mouse proceeded through the maze to make a nose-poke response in an arm, at which time the LEDs turned off. Correct responses (matching the side of the LED light cue) were rewarded with 15µL of 10% sucrose solution, while incorrect responses were indicated by 1s of white noise (~68dB). The LED light cue side (left/right) was determined randomly, unless the mouse poked the same arm three trials in a row, in which case the LED light cue side was set to the opposite side. Testing continued for 30 minutes or until the mouse completed 100 trials, whichever occurred first. Daily behavioral testing began when the mouse was approximately 80 days of age and continued through day 114. Test sessions with fewer than 25 trials were excluded from analysis. Ten minutes of dim-light open field recordings were acquired for each mouse on the day before testing began. A total of 217 WT and 199 MU test sessions were included in analysis.

Prior to testing, mice completed three training stages. In stage 1, the goal was to learn the maze layout. Mice could nose-poke any hole for a reward of 10µL of 10% sucrose, and the training stage was passed when mice completed 2 consecutive training days of at least 30 pokes in 30min with more than one poke per hole. In stage 2, a shaping stage, the goal was to learn the stem-arm response pattern. Mice poked the stem to start each trial for a reward of 5µL of 10% sucrose, and next had to poke in an arm to receive 15µL of 10% sucrose. Incorrect pokes were unrewarded. This training stage was passed when mice completed 2 consecutive days of at least 20 correct trials in 30min. In stage 3, the goal was to learn the cue-response association. Mice poked the stem to start each trial for a reward of 5µL of 10% sucrose, and either the right or left LED light cues turned on. To get rewarded, mice had to poke in the arm indicated by the LED for 15µL of 10% sucrose. Incorrect pokes were unrewarded. This training stage was passed when mice completed 2 consecutive days of at least 25 trials in 30min. To encourage responding in both arms during the task and in training stages 2 and 3, if the same arm was poked on three consecutive trials, the opposite arm was required for reward on the next trial.

**Delayed Sample-to-Match T-Maze Task**

This task was the same as the sample-to-match task, with the addition of a difficulty modulation in the form of a working memory delay. The mouse poked the stem to start each trial, getting rewarded 5µL of 10% sucrose solution. Then the gate closed, and mice were shown either the right or left LED light cue. The mouse had to poke the poke hole near the gate that matched the side of the light cue (cue poke hole), at which time the LED turned off, and a delay commenced. After the delay, the gate opened, and from here, each trial was identical to the sample-to-match task. Mice performed 75 trials per session, divided into 3 blocks of 25 trials each. Each block of 25 consisted of five 3-second trials, followed by five1.5-second trials, five 8-second trials, five half-second trials, and five 5-second trials. A total of 153 test sessions were included in analysis.

The first three training stages for this task were identical to the training stages for the sample-to-match task. In stage 4, mice learned to open the gate by poking the poke holes near the gate corresponding to the LED cute (in the final task, this is what commences the delay period). The mouse poked the stem to start each trial, getting rewarded 5µL of 10% sucrose solution and triggering the gates to close. Mice were shown either the right or left LED light cue and had to poke the hole near the gate that matched the side of the light cue to open the gates. Mice then had to respond in the arm corresponding to the LED cue to receive 15µL of 10% sucrose solution, with incorrect responses indicated by the white noise as in the task. This stage was passed when mice completed two consecutive sessions with 65% task accuracy and at least 25 trials in 30min. In stage 5, mice were introduced to the delay. Stage 5 is identical to stage 4, except now when the mouse pokes cue poke hole, the gates open after a delay, rather than immediately. The delay pseudo-alternated between a half-second delay and a dynamically determined variable delay. The variable delay was initially 1s, and increased by 200ms for every incorrect trial, and decreased by 500ms for every incorrect trial to a minimum of 1s. This stage was passed when mice completed 2 consecutive days reaching a variable delay of 3s and completing at least 25 trials in 45min. To encourage responding in both arms during the task and in training stages 2 onward, if the same arm was poked on three consecutive trials, the opposite arm was required for reward on the next trial.

**Neural Electrophysiological Data Acquisition & Video Recording**

Neurophysiological recordings were acquired during behavioral testing with the Cerebus acquisition system (Blackrock Microsystems, Inc., UT). Extracellular neuronal activity was sampled at 30kHz, high-pass filtered at 500Hz, and sorted online. Online sorting consisted of referencing data against a wire within the same brain area that did not exhibit a signal-to-noise ratio greater than 3:1. After full recording, cells were sorted again using an offline automated sorting algorithm (Plexon Inc., Dallas, TX) and additionally verified by investigators to confirm the quality and identification of recorded cells. Local field potentials (LFPs) were sampled and stored at 1kHz, bandpass filtered at 0.5-250Hz. All neurophysiological data were referenced to a ground wire connecting the ground screws above cerebellum and anterior cranium.

Video recordings were acquired in real time using NeuroMotive and synchronized with neurophysiological data. Mouse position was tracked offline using a mean-shift algorithm as implemented via Python OpenCV (https://docs.opencv.org/3.4.0/index.html). Position was extracted from each video. In order to ensure that all position and velocity data were in standard space, the boundaries of the T-maze were traced in each video recording. A "standard T" was defined as the mean of all of these boundaries, and a projective linear transformation was used to transform each recording to this standard space, and for instantaneous velocity, to further convert pixel measurements to meters. The velocity time series was smoothed with a 250ms sliding window.

**Data Analysis: Task Epochs**

During performance of the T-maze, anticipation during the arm runs was defined as the period between the gates opening and the mouse's nose poke response. Anticipation during the stem runs was defined as the period between the mouse's arm nose poke response and the mouse's subsequent stem nose poke response. Goal proximity was operationalized as the percent goal progress along the mouse's traveled path. Note that in order to minimize the effect of the nose poke responses on anticipatory network investigation, the anticipatory period was constrained to 5-95% goal progress. Activity just before goal was operationalized as mean activity at 90-95% goal progress, and in the delayed sample-to-match task, the period just after the gate opened was operationalized as mean activity at 0-5% goal progress.

**Data Analysis: Mixed Effects General Linear Model**

Since multiple test sessions were acquired per mouse, all test statistics were calculated per test session. For all analyses that included multiple test days per mouse (e.g., arm vs. stem component activity, correlation of component with velocity, correlation of component or other neural measures with goal progress, behavioral accuracy, etc), group statistics accounted for repeated measures via mixed effects general linear models with random intercept for mouse. Results were subject to Bonferroni correction for multiple comparisons.

**Independent Components Analysis (ICA) of Power Spectrograms**

Functional networks were operationalized as groups of regional frequency bands whose power were temporally correlated. ICA, a data-driven machine learning approach, on power spectrograms was used to identify putative functional networks. ICA has been applied to LFP power spectrograms and has also been used to identify task-based as well as resting state networks in human functional magnetic resonance imaging data[5-7]. LFP recordings were high-pass filtered over 1Hz, notch-filtered at 60Hz and all harmonics to remove electricity artifact. Saturated timepoints, defined as an amplitude of six-fold standard deviation or greater were also removed. Power spectrograms for 4-160Hz were generated from the LFP recordings, using short-time Fourier transform as implemented in Matlab 2016a (Mathworks, Inc.), calculated over a sliding window of 500ms with a 10ms step. Power spectrograms were averaged across electrodes in each region.

Training data for ICA consisted of normalized 10-minute power spectrogram segments randomly selected out of each of 10 randomly chosen days from each of the 7 WT mice during the sample-to-match task, downsampled by a factor of 10 to an effective sampling rate of 10Hz. ICA, as implemented via the FastICA algorithm in scikit-learn version 0.19.1, a Python machine learning library (http://scikit-learn.org/stable/index.html), was used to decompose the data into independent non-Gaussian components. Three to 942 components were tested, where the maximum number meant each component consisted of a single 1Hz frequency band from a single region. To determine the number of components to use, a validation data set, which consisted of normalized 10-minute power spectrogram segments randomly selected from each of previously unused 5 randomly chosen days from the 7 WT mice, were projected into the identified components to assess the proportion of variance explained by the components. To balance complexity (i.e., prioritizing the % variance explained) with parsimony (i.e., prioritizing a small number of components), the number of components was chosen using the elbow method, which identifies the point at which adding more components results in less gain in explained variance. For all analysis of network activity, power spectrograms were projected into the ICA-learned networks via the unmixing matrices, or loading matrices, at the same temporal resolution as the power spectrograms, i.e., 100Hz.

**LFP Directionality Analysis**

Directionality was inferred from LFP pairs based on cross-correlation of instantaneous phases of oscillations at varying temporal lags[8,9]. For each recording, two electrodes were chosen at random for each region and LFPs for each electrode filtered using Butterworth bandpass filters to isolate the LFP oscillations within the frequency of interest. Instantaneous phase of the filtered LFPs was determined using the Hilbert transform, and instantaneous phase offset was calculated as the difference of the phase time series, and mean resultant length (MRL) of the distance was calculated, corresponding to the deviation from circular uniformity (where 0 represents no net deviation from uniformity and 1 represents complete deviation from uniformity at a single angle/phase)[9,10]. This was recalculated at each temporal offset ranging from -100ms to 100ms in 10ms increments, and the temporal offset that yielded the highest MRL was determined to be the temporal offset of optimal phase coupling, allowing for inference of signal direction flow. A frequency and brain area pair were deemed to exhibit significant directionality if the group mean (mixed effects general linear model with random intercept for mouse), was significantly different from 0. Since we were mainly interested in drivers of the network when it was being engaged, we isolated segments corresponding to high component activity (>median) with enough timepoints (200ms) for this analysis.

**LFP Coherence**

For each pair of regions, two electrodes were selected at random, and cross-area coherence was calculated from each microwire LFP pair using magnitude-squared coherence, as implemented via *mscohere* in Matlab2016a (MathWorks, Inc.). Coherence was calculated for 500ms sliding windows, with a 100ms step. These calculated coherence values were averaged across microware pairs. Coherence was averaged within frequency bands of interest.

**Correlation of Cell Firing with Component Activity**

We employed a conservative approach to ensure that cells were not double counted in our analysis. Specifically, we restricted analysis of cells to one test session per microwire implant (i.e., channel). We chose the test session with the most recorded cells for a given channel, and if there were multiple sessions with the same number of recorded units in a given channel, we randomly chose one. To test whether cell firing rate correlated with network activity above and beyond goal progress, cell spiking was first transformed into a time series matching the LFP sampling rate (100Hz). We then isolated the anticipatory period from each trial (i.e., 5-95% goal progress), and binned it into bins of 5%, such that each anticipatory period had 18 bins. Firing rate and component activity were averaged for each bin. The bins were concatenated across trials and Pearson correlation coefficient calculated. To generate a bootstrapped null distribution, the trial labels were shuffled for the cell spikes 10000 times. If cell firing were merely a function of goal progress, shuffling trial labels would preserve this correlation, since they would still be lined up with the same goal progress percentage. Cells whose Pearson correlation coefficient was less than the 2.5th percentile or greater than the 97.5th percentile of the bootstrapped null distribution were considered to be significantly correlate with the network above and beyond goal progress.
